# Spleen gene expression is associated with mercury content in three-spined stickleback populations

**DOI:** 10.1101/2024.04.29.591498

**Authors:** Brijesh S. Yadav, Fabien C. Lamaze, Aruna M. Shankregowda, Vyshal Delahaut, Federico C. F. Calboli, Deepti M. Patel, Marijn Kuizenga, Lieven Bervoets, Filip A.M. Volckaert, Gudrun De Boeck, Joost A.M. Raeymaekers

## Abstract

Mercury can be very toxic at low environmental concentrations by impairing immunological, neurological, and other vital pathways in humans and animals. Aquatic ecosystems are heavily impacted by mercury pollution, with evidence of biomagnification through the food web. We examined the effect of mercury toxicity on the spleen, one of the primary immune organs in fish, in natural populations of the three-spined stickleback (*Gasterosteus aculeatus* Linnaeus, 1758). Our aim was to better understand adaptation to high mercury environments by investigating transcriptomic changes in the spleen. Three stickleback populations with mean Hg muscle concentrations above and three populations with mean Hg muscle concentrations below the European Biota Quality Standard of 20 ng/g wet weight were selected from the Scheldt and Meuse basin in Belgium. We then conducted RNA sequencing of the spleen tissue of 22 females from these populations. We identified 136 differentially expressed genes between individuals from populations with high and low mean mercury content. The 129 genes that were upregulated were related to the neurological system, immunological activity, hormonal regulation, and inorganic cation transporter activity. Seven genes were downregulated and were all involved in pre-mRNA splicing. The results are indicative of our ability to detect molecular alterations in natural populations that exceed an important environmental quality standard. This allows us to assess the biological relevance of such standards, offering an opportunity to better describe and manage mercury-associated environmental health risks in aquatic populations.

## Introduction

Chemical pollution has rapidly changed aquatic ecosystems worldwide (Hooper et al., 2012). Metals, including mercury (Hg), are of particular concern because of their omnipresence and availability for uptake in biota, in addition to their high potential for interference with the functioning of vital cellular components (Okereafor et al., 2020). Mercury is released into the atmosphere from natural and anthropogenic sources and deposited globally. It may accumulate in the food chain in its organic form, methylmercury (MeHg). Once accumulated, mercury may exert toxic effects on various biological pathways and alter immune responses, neuronal signalling, and osmoregulation, leading to organ failure (Yang et al.,2020, Balali-Mood et al., 2021). In fish, mercury concentrates in organs such as the liver, muscle tissue, gills, kidney and brain through exposure to contaminated water or food (Morcillo et al., 2017, Zulkipli et al., 2021).

Some fish populations thrive at sites with contaminant concentrations that elicit toxic responses in naive populations (Durrant et al., 2011). Such resistance may involve intragenerational physiological acclimation and intergenerational genetic adaptation (Reid et al., 2016). Individual organisms may cope with toxicity through regulatory or epigenetic modes of adaptation (Hu & Barrett 2017). Yet, adaptation does not always occur where expected, as populations are often rapidly extirpated at contaminated sites (Coffin et al., 2022). The study of adaptation to contaminants is thus crucial for adequate environmental risk assessment. Many studies on experimental as well as wild populations have improved our understanding of how mercuric toxicity disrupts metabolic, endocrine and cellular pathways, and have provided a basis for the development of biomarkers for mercury pollution (Driscoll et al., 2013, Branco et al., 2017, Vasconcellos et al., 2021, Olsvik et al., 2021, Trivedi et al., 2022). In contrast, relatively few studies have investigated how genetic adaptation shapes the biological responses to mercury pollution and, therefore, how populations under pollution stress ultimately manage to persist (Belfiore and Anderson, 2001, Budnik and Casteleyn, 2019, Yang et al., 2020). Such studies require the comparison of these responses between natural populations with a history of high versus low mercury contamination.

In this study, we investigate the effects of MeHg in natural three-spined stickleback (*Gasterosteus aculeatus* Linnaeus, 1758; Gasterosteidae) populations in Flanders (Belgium). The three-spined stickleback is increasingly used as a model species in ecotoxicological studies (Katsiadaki et al., 2002, Willacker et al., 2013, Webster et al., 2017, Calboli et al., 2021). Flanders has experienced a significant degree of metal pollution from industrial activities since the 19th century, with very local though scattered signatures of mercury contamination in soils and sediments (Tack et al., 2005; Van Steertegem, 2011, Brosens et al., 2015). Accordingly, mercury concentrations in the muscle tissue of stickleback populations are highly variable, ranging from 21.5 to 327 ng/g dry weight (Calboli et al., 2021). Calboli et al. (2021) further revealed that Hg accumulation is associated with specific genomic regions, suggesting a genetic basis for adaptation to mercury-polluted environments in three-spined sticklebacks in Flanders.

Here, we further investigate the effects of mercury on these three-spined stickleback populations by means of a gene expression study in the spleen tissue. The analysis of genome-wide variation in gene expression is particularly relevant, as it facilitates the interpretation of an organism’s basic molecular mechanisms and phenotypic plasticity in response to toxicity (Reid et al., 2016). The spleen, crucial in fish immune defense and blood filtration, is known to be affected by mercury pollution (Tjahjaningsih et. al., 2017). For instance, exposure of carp to mercury chloride led to a notable increase in macrophage accumulation in the spleen tissue, suggesting that spleen tissue may serve as a sensitive bioindicator for environmental pollution (Kaewamatawong et.al., 2013; Tjahjaningsih et. al., 2017). Mercury exposure may also impair the functioning of the spleen’s white pulp tissue (Roales and Perlmutter 1980). However, the details of these impairments are not fully understood. We therefore investigated the spleen’s functions at the transcriptome level to better understand adaptation to high mercury environments. Specifically, we hypothesise that the spleen’s immune functions are linked with pathways underlying mercury-associated stress responses. By comparing transcriptomic profiles among three-spined sticklebacks from populations with high and low mercury accumulation, we explore molecular targets and identify genes and pathways of environmental mercury exposure. By focusing on populations with mean Hg concentration above and below the European Biota Quality Standard of 20 ng/g wet weight, we aim to better understand the biological relevance of such standards.

## Material and Methods

### Site selection, sampling and sample preparation

The biological materials for this study were collected in parallel with the study by Calboli et al. (2021), who described the variation in mercury concentrations in the muscle tissue among 523 sticklebacks from 21 locations and performed a genome-wide association study on the accumulated levels of mercury in these individuals, based on 28,450 SNPs. The 21 populations originated from rivers and streams within three drainage basins in Flanders (Belgium), namely the river Meuse basin, the eastern basin of the Scheldt River (Scheldt-E), and the western basin of the Scheldt River (Scheldt-W) (Figure 1A).

**Figure 1.**
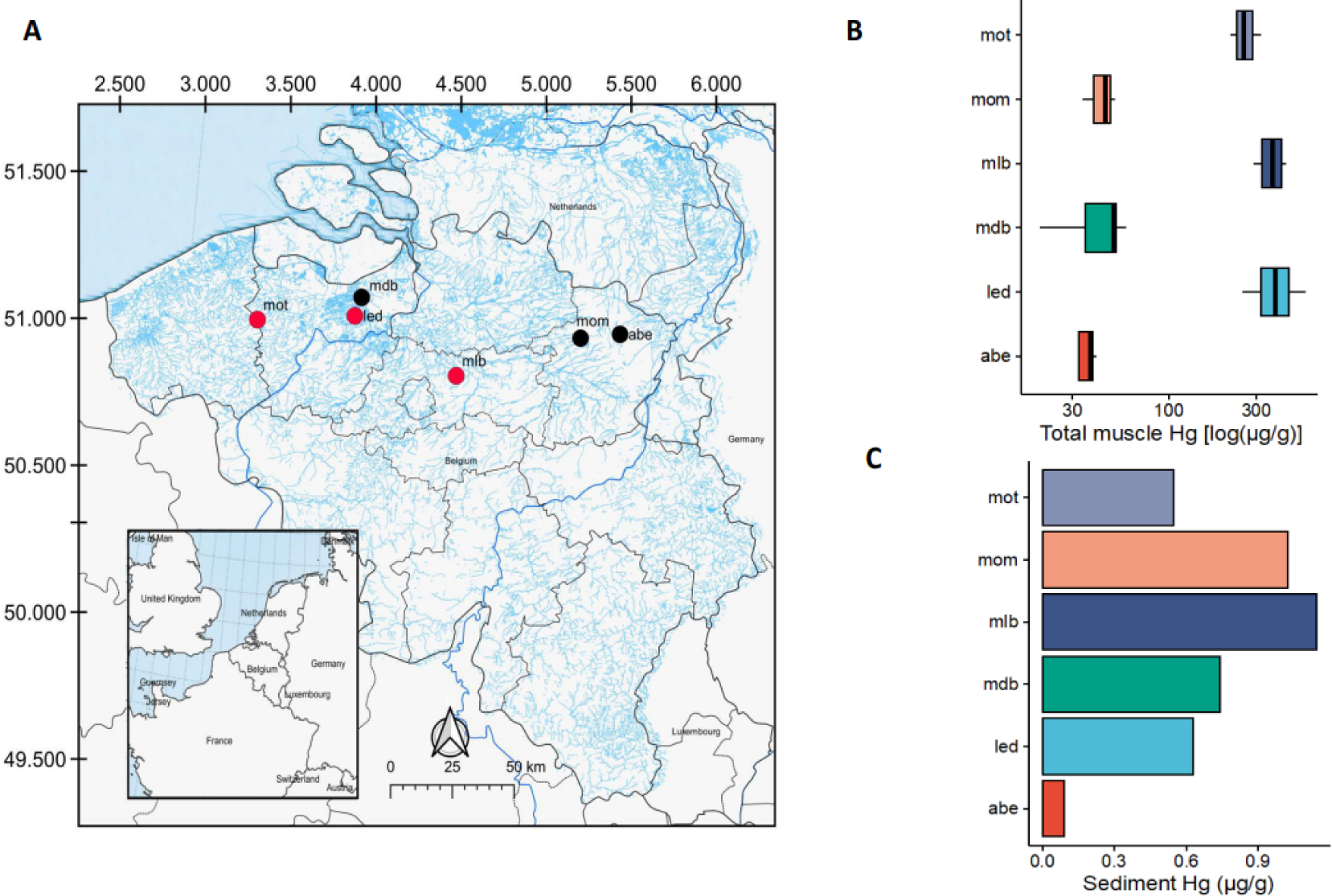
(A) Map of the sampling region with indication of the locations of the 21 study populations. Locations in red and grey mark the subset of populations with high and low mean mercury concentrations in muscle tissue, respectively. (B) Log-transformed total muscle Hg concentrations across the six focal stickleback populations. (C) Hg concentration in the sediment of the six focal locations.

An overview of sampling sites, sample collection, sample preparation, sex determination, and measurement of mercury content in muscle tissue and in the sediment is available in Calboli et al. (2021). Here, we provide a summary of these methods, and describe additional procedures relevant for our study. About 25 individual sticklebacks per location were collected using dip nets. Fish were then transferred to an oxygenated bucket filled with 25L of water collected on site, and transported to the University of Antwerp (Antwerp, Belgium).

The fish were kept overnight and were then euthanized using an overdose of buffered Tricaine mesylate (MS-222) according to the guidelines of the Ethical Commission for Animal Experiments of the KU Leuven. Individual fish were weighed (W_B_; mg) and measured for standard length (SL; mm). Fish were dissected on a glass plate kept on ice to maximize tissue preservation. In case the gonads were mature, the sex of the sample was recorded. The liver was extracted and weighed (W_L_; mg). The spleen (range: 0.1-3.2 mg) was collected and stored in RNA later (Thermo Fisher Scientific – Invitrogen, Waltham, MA, United States). For mercury analysis, muscle tissue on both flanks was dissected and snap-frozen in liquid nitrogen and temporarily stored at −80°C. For the measurement of mercury content in muscle tissue as well as in the sediment, we refer to the procedures described in Calboli et al. (2021).

For the present study, a subset of six populations was selected for the analysis of gene expression. This subset included three populations with low mean mercury concentrations in muscle tissue (Figure 1A; Abeek [*abe*]: 22.0 ± 13.9 ng.g^−1^; Molenaarsdreefbeek [*mdb*]: 49.9 ± 20.9 ng.g^−1^; Mombeek [*mom*]: 102 ± 128 ng.g^−1^), and three populations with high mean mercury concentrations in muscle tissue (Lede [*led*]: 322 ± 180 ng.g^−1^; Molenbeek [*mlb*]: 326 ± 77.8 ng.g^−1^; Motebeek [*mot*]: 224 ± 70.0 ng.g^−1^) (all values measured as dry weight concentrations). We further refer to the two sets of populations as the “high Hg” and “low Hg” group. Assuming a typical moisture content of 80 % in fish tissue, these mean dry weight concentrations correspond to 4 to 65 ng/g wet weight, and all three populations of the high Hg group exceed the European Biota Quality Standard of 20 ng/g wet weight (Ribeiro et al., 2015, <u>European Commission, 2013</u>).

Mercury concentrations of the six focal populations differed significantly between the high and low Hg groups after correction for standard length (Figure 1B; F_5, 141_ = 59.1; p-value < 0.0001). Yet, there was no correlation between the mean mercury concentration in muscle tissue and the mercury concentration measured in the sediment (Figure 1C; Spearman rho = 0.54, p-value = 0.30), as already indicated for the complete set of populations (Calboli et al. 2021). Furthermore, with three locations from Scheldt-E (*mot*, *led* and *mdb*), two locations from Scheldt-W (*mlb* and *mom*), and one location from the Meuse basin (*abe*), we ensured that there was no geographic clustering between populations of the high Hg and low Hg group.

### RNA extraction, library preparation, sequencing and bioinformatic analysis

#### RNA extraction

Total RNA was extracted from the spleen tissue of 24 females from the six focal populations, with three to five individuals per location. The individual whole spleen tissues were thoroughly homogenised with 500 ml of TRI reagent (Zymoresearch, USA) using 1.4 mm ceramic beads (Genaxxon Bioscience, Germany) at 6,500 rpm for 2 × 20 s in a Precellys 24 homogenizer (Bertin Instruments, Montigny-le-Bretonneux, France). The RNA was extracted from the supernatant of the tissue homogenate using Direct-zol™ RNA MiniPrep (Zymoresearch, USA), following the manufacturer’s instructions. RNA purity and quality were assessed using NanoDrop™L 1,000 (Thermo Fisher Scientific) and TapeStation (Agilent Technologies, USA). The RNA quantification was performed with a Qubit fluorometer (Thermo Fisher Scientific). The RNA integrity number (RIN value) was > 8 in all samples.

#### Library preparation and sequencing

Twenty-four individual RNA libraries were constructed with the NEBNext® Ultra™ II Directional RNA Library preparation kit (NE Biolabs, USA) and poly(A) mRNA magnetic isolation module (NE Biolabs) with the given procedure. The library preparation started with 1 µg of total RNA, and after Poly(A) enrichment, mRNA was fragmented (the samples were heated at 65 °C for 5 min) to obtain 100–200 bp fragments. Then, the first and second strands of cDNA were synthesised and purified. The next steps, adaptor ligation and barcoding, were performed with NEBNext® Multiplex Oligos (NE Biolabs), followed by PCR enrichment with 9 cycles. Amplified libraries were purified utilising Mag-Bind TotalPure NGS (Omega Bio-tek, United States). In the last step, libraries were pooled according to barcodes and loaded at 1.4 pM on the Illumina NextSeq 500 sequencer (Illumina) with the NextSeq 500/550 High Output Kit (v2.5, 75 cycles) for 75 bp single-end sequencing at the genomics platform of Nord University (Bodø, Norway). On average, about 41 million single-end raw reads were obtained for each sample. Yet, two samples from location *mot* were of insufficient quality after library preparation or sequencing and were therefore excluded from further analyses.

#### Bioinformatics analysis

The adaptor sequences were trimmed using the fastp software (Chen et al., 2018) with default parameters. Phred quality score (Q ≥ 30) was used for filtering raw reads, and the quality of the filtered reads was checked using the FastQC software. Cleaned reads were aligned to the reference genome (Peichel et al., 2020, Nath et al., 2021) of the three-spined stickleback (stickleback_v5_assembly.fa.gz, retrieved directly from the Stickleback Genome Browser hosted by the University of Georgia) using HISAT2, version 2.2.1 (Kim et al., 2019). Subsequently, the feature Counts tool (<u>Liao, Smyth and Shi, 2014</u>) was used to find the read counts that belong to each transcript using the transcriptome (stickleback_v4_ensembl_lifted.gtf). Genes were filtered to read counts > 10 in each sample prior to further analysis.

### Data analysis

#### Phenotypic traits

To assess the relative condition of the individual fish of the six focal populations, two condition factors were used: (1) Fulton’s condition index (CI) was calculated as 1000 × (W_B_/SL^3^); and (2) the hepato-somatic index (HSI; a proxy of energy reserves) was calculated as HSI = (W_L_/W_B_) × 100, where W_L_ and W_B_ represent the wet liver weight and wet body weight, respectively. Body size (measured as SL), body weight, and relative condition (CI and HSI) were compared using a nested ANOVA with mercury group (high Hg vs. low Hg) as a fixed factor and sampling location (nested in mercury group) as a random factor.

#### Population genetic structure

To correct differences in gene expression for genetic background (see below), we quantified the population genetic structure of the 21 populations using a Principal Component Analysis (PCA). Data from 512 individuals and 28,450 SNPs were retrieved from Calboli et al. (2021). SNPs identified to be under selection, associated with variation in Hg concentration in muscle tissue (c.f. Calboli et al., 2021), linked to sex chromosome XIX (<u>Peichel et al., 2004</u>), or having any missing values across the 512 individuals were excluded from the analysis. The *PCA()* function from the FactoMineR (version 2.4) R package was used to perform the PCA analysis.

#### Differential gene expression

We compared gene expression between individuals from the high mercury environment with individuals from the low mercury environment. A negative binomial distribution model was fitted with six predictors, including one covariate associated with high vs. low Hg, and five covariates associated with genetic background. The genetic background predictors consisted of the five first principal components of the PCA performed to assess population genetic structure (see above and Supplementary Figure 1). The projection was scaled to avoid overdispersion with the *scale()* function. The DESeq2 package (version 4.1) (Love, Huber and Anders, 2014) was used to fit an additive model (AM) as G_i_ ∼ M_i_ + PC1_i_ + PC2_i_ + PC3_i_ + PC4_i_ + PC5_i_, with G_i_ being the gene investigated; M_i_ the Hg group (high vs. low Hg in muscle tissue), and PC1_i_ to PC5_i_ the loadings of each individual on each principal component. Differential gene expression was assessed with *lfcShrink().* Genes with a cut-off value of |Log2(fold change)| ≥ 1 and an adjusted s-value of < 0.1 (local false sign rate) were considered differentially expressed. To visualise the differentially expressed genes, we used an MA plot (log2 fold change versus log2 expression). Finally, to visualise the expression differences between the 22 individuals, we used the *PCA()* function from the FactoMineR package.

#### Enrichment analysis, Netwok Analysis and Pathways

we used the differentially expressed genes for gene ontology analysis using g:Profiler’s g:GOSt tool (https://biit.cs.ut.ee/gprofiler/gost), targeting the Ensembl three-spined stickleback database. We focused on the list of differentially expressed genes, setting our statistical domain against all known genes. The significance threshold was established using g:SCS at a user-defined level of 0.05, and for identification purposes, numeric IDs were treated as ENTREZGENE_ACC.

We used Cytoscape (version 3.9.1) to make a canonical protein-protein interaction (PPI) network by utilising the STRING web-based database (version 11.5) to evaluate the relationship among the 129 upregulated genes. In this analysis, we used the background database from zebrafish *Danio rerio.* We performed pathway enrichment analysis by applying Gene Set Enrichment Analysis (GSEA) with 1000 permutations and FDR parameters set to less than 0.05. In order to identify gene (co)regulatory dysregulation, we carried a Delta network analysis by first calculating the Pearson correlation among differentially expressed genes within the high and the low mercury group. Then we subtracted the high and the low mercury correlation matrices to compute the delta matrix. Using the *r.test()* from the psych package (version 2.2.9), we computed the p-values associated with the dysregulation of the gene network (delta net) and used an FDR of < 0.01 to consider the hub (i.e., a node interconnecting a gene with other genes) to be differentially regulated. For presentation, only the dysregulatory hubs with ten connections or more were included.

## Results

### Population characteristics and genetic structure

Neither body size, body weight, condition index, nor HSI differed between the populations of the high Hg and the low Hg group (all p-values > 0.05) (Figure 2A-D). As previously observed by Calboli et al. (2021), the population genetic structure of the 21 populations reflected watershed and sampling location, rather than Hg levels (Figure 2E). The 22 individuals used for gene expression analyses also clustered by population, rather than by high vs. low Hg group (Figure 2F).

**Figure 2.**
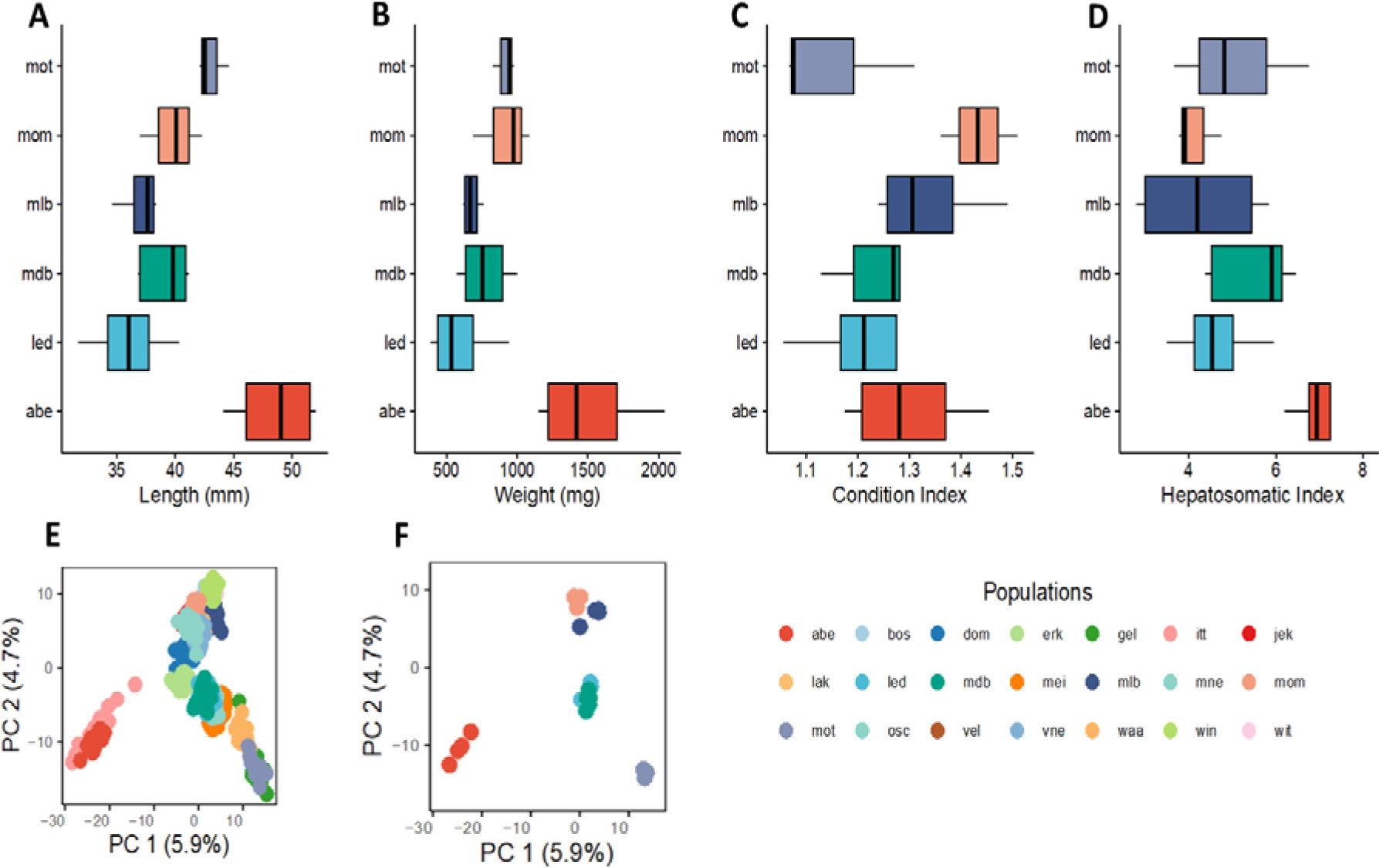
Population characteristics and genetic structure. (A) Standard length, (B) body weight, (C) condition index, (D) hepatosomatic index. (E-F) SNP-based PCA plot representing the genetic relationship among (E) all 21 populations and (F) the 22 individuals from the six focal populations selected for gene expression analysis.

### RNA sequencing output

About 922 million raw reads were obtained from the 22 spleen samples (Supplementary Table 1). About 2 % of reads were lost during filtering for read quality, resulting in 904 million reads. The average percentage of bases with a Phred quality score of Q ≥ 30 was 95.02 %. The average mapping percentage of filtered reads and GC content was 90.7 % and 51.3 %, respectively.

### Differential gene expression reveals common signatures

A total of 136 genes were differentially expressed between the individuals from the high vs. low Hg group. The MA plot (Supplementary Figure 2 and Supplementary Table 2) indicated that a total of 129 genes were upregulated in the high Hg group, while seven genes were downregulated. A heatmap of the 136 genes combined with hierarchical clustering revealed one cluster comprising two individuals of the high Hg group (population led; Led03_H and Led04_H), and two mixed clusters with individuals of both groups (Figure 3A).

**Figure 3.**
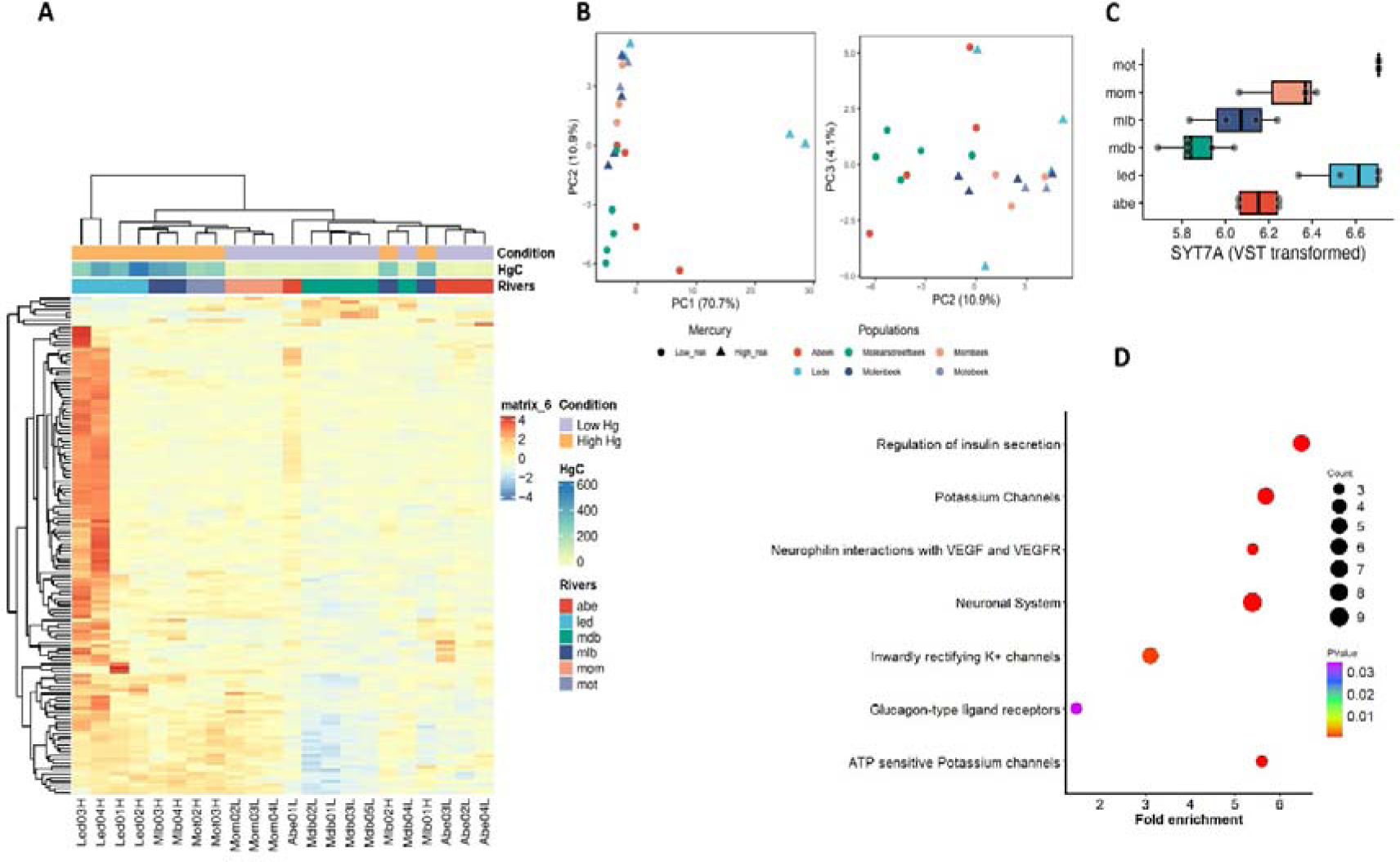
Differentially expressed genes are associated with the neuronal and transport system. (A) Hierarchical clustering of transcriptome-based differences between individuals of the high Hg and low Hg group. The heatmap was built with DeSeq2 based on variance normalised read counts. The heatmap shows the differentially expressed genes with an adjusted s-value below 0.1 and |Log2 fold change| ≥1. (B) PCA plots (PC1 vs. PC2 and PC2 vs. PC3) visualising the variation in expression for the 136 genes that are differentially expressed between individuals from the two Hg groups. (C) Boxplots of differential expression by location for SYT7, a gene involved in neuronal functions. (D) Pathway enrichment analysis of the 129 upregulated genes, as identified through the STRING database and analysed in Cytoscape, using FDR values for significance determination. The enrichment data, originating from Reactome Pathways, involved biological functions. The fold enrichment of pathways is depicted on a log-transformed scale. Circles within the figure vary in size proportionally to the count of genes associated with each pathway. The color intensity of each circle represents the associated p-value.

A PCA plot based on the set of 136 differentially expressed genes separated two individuals from population led (Led03_H and Led04_H) from all other individuals along PC1, while PC2 showed a cline from mostly low Hg individuals (negative PC2 values) to more high Hg individuals (positive PC2 values) (Figure 3B). Gene expression levels for selected genes with neuronal functions (SYT7A in Figure 3C; RIMS2 and CHRNB2 in Supplementary Figure 3) were consistently high for population *led*, but were more variable for the two other populations of the high Hg group, *mlb* and *mot*.

### Differential expression levels between the high and low Hg group reveal an activation of neuronal-like functional pathways and rewiring of gene regulatory networks

Enrichment analysis on 129 upregulated genes using Reactome pathways revealed that the neuronal function including potation channels regulation are significantly enriched, representing 4 out of 7 enriched pathways (Figure 3D). This is supported by a hierarchical enrichment term examination using a GO-terms-based analysis (Supplementary Figure 4). We identified 3/3 biological processes associated with the regulation of localization and transport function activity. At the cellular component level, 6/6 terms were related to cell membrane and secretory activities. Finally, at the molecular functions level 8/12 terms were involved in channel transport regulation. Further at the gene level, by annotating a canonical protein-protein interaction network with Cytoscape for biological networks we found out of 79 gene proteins that 33 (51 %) were associated with the neuronal systems (axon and synapse), 13 (20 %) with plasma membrane proteins, 12 (18 %) with inorganic transporter activity, 11 with secretory vesicles, 8 (12 %) with functions related to potassium, and 4 functions related to calcium ion transport (Figure 4A).

**Figure 4.**
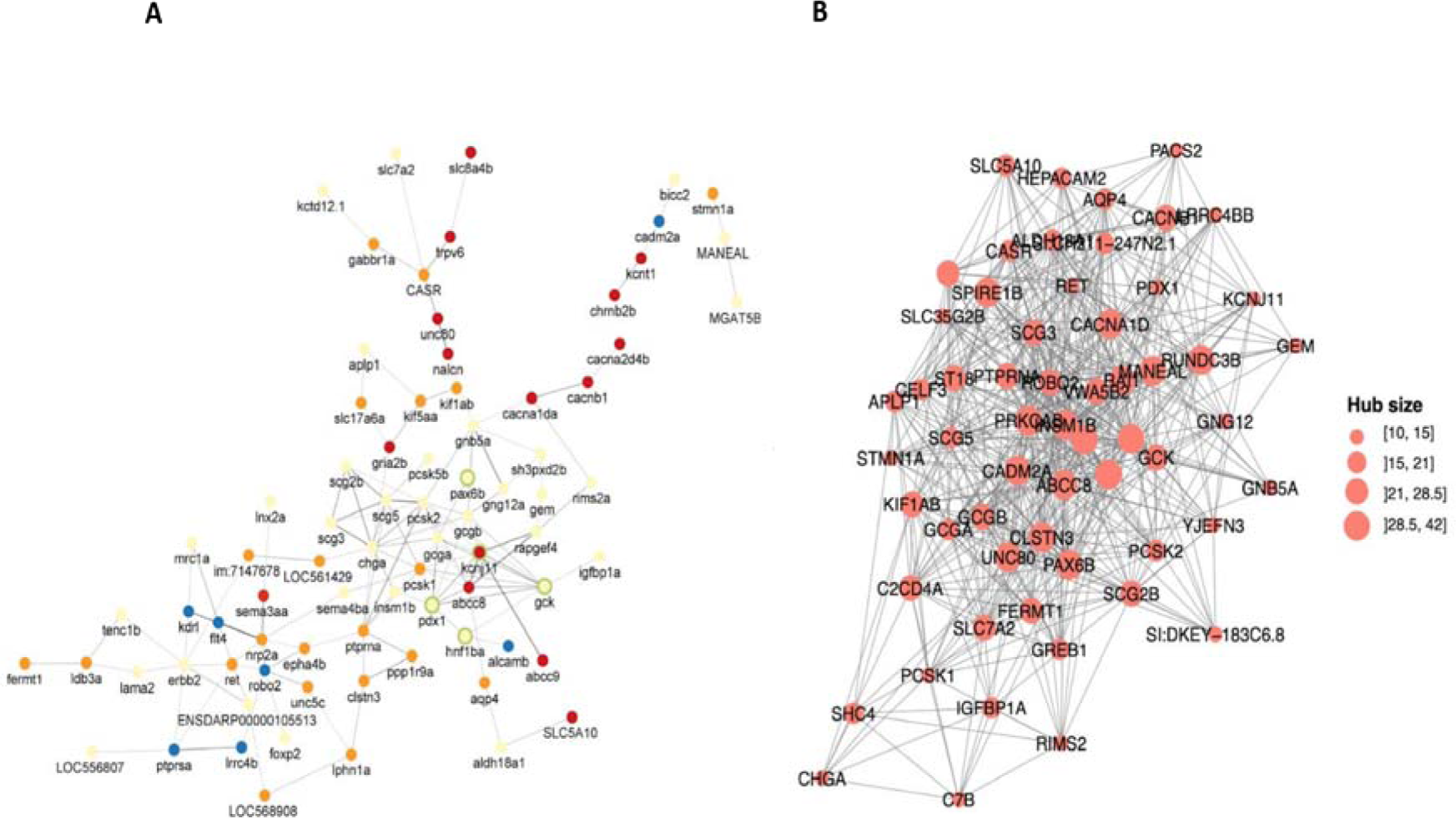
Regulatory networks of differentially expressed genes. (A) Protein-protein interaction (PPI) network based on the 129 upregulated genes. The PPI enrichment score was 1.0 E-16, with a confidence score of 0.5 (fdr < 0.05), underlying 206 functional associations between 65 genes (nodes). These nodes are involved in the nervous system (yellow), plasma membrane (orange), Immunoglobulin (blue), and inorganic transporter activity (red). Nodes with a green circle mark genes with a pancreatic b cell function. The confidence score of each interaction is proportional to the edge thickness and opacity. (B) Delta network, depicting the significantly dysregulated hubs genes between individuals from the high Hg and low Hg group (FDR < 0.01). Only the significant hubs of at least ten interactions are represented.

The canonical protein-protein network of upregulated genes in the high Hg environment reveals that the protein hub associated with the nervous system and plasma membrane are central in the network with maximal edges interactions, while other functions are peripheral (Figure 4A). Furthermore, we tested which gene and gene interactions are more likely to be dysregulated by Hg intoxication by using a delta analysis between the gene interaction networks of the high Hg vs. low Hg group by subtracting the corresponding gene expression correlation matrices (Supplementary Figure 5). This delta network indicated significant regulatory rewiring of central gene hubs (Figure 4B; Supplementary Figure 5). Genes with extreme breakdown of coordinated expression were associated with the nervous system (e.g., GCGA, GCGB, PDX1) and plasma membrane (e.g., RET) (Figure 4A).

## Discussion

We compared the transcriptional responses between three-spined stickleback populations with Hg levels above (high Hg group) and below (low Hg group) the European Biota Quality Standard of 20 ng/g wet weight. We hypothesised that the toxic effects of Hg above this standard affect the immunity of individuals assessed by the transcriptional repertoire of the spleen, an important lymphoid organ. We found 136 differentially expressed genes between the individuals from the two groups, not only confirming our ability to associate transcriptomic alterations with metal exposure in natural vertebrate populations (Maes et al. 2013), but also indicating that there is a measurable biological response when an important environmental quality standard is exceeded. Furthermore, we showed that these alterations involved not only genes associated with the cell membrane’s physiological systems and various immunological functions but also the neurological system. Indeed, both the analysis of differential gene expression and the enrichment and network analysis revealed functions associated with neuron projection, axon activity, and synapse functioning. This suggests that gene expression in the spleen involving cell communication, calcium transport, and neurotransmitter release are interconnected, and the homeostatic regulation of the neuronal-immune crosstalk is affected upon mercury intoxication. Finally, we found that gene expression was rather variable within the high Hg group, suggesting that exposure to high Hg can lead to diverse biological signals.

The detected differences in gene expression between the high Hg and low Hg group represent contrasts between natural populations, and therefore do not allow us to infer causality. By statistically correcting for neutral genetic background, we ensured that the results do not merely represent random population-level differences that evolved over time. Yet, other potential differences were not accounted for. Specifically, although the six populations originate from fairly similar lowland rivers and streams, the difference in Hg content between the two groups may reflect environmental features that could also affect gene expression patterns. Nevertheless, our findings are in line with a diverse set of experimental and field-based studies highlighting the complex and varied impacts of Hg exposure in fish. In nature, it has been observed that fish living in high Hg environments have altered gene expression profiles in detoxification and oxidative stress pathways compared to those living in low Hg environments (Carvalho, 1993, Brady et al., 2017, Adrian-Kalchhauser et al., 2020, Burke et al., 2020, Olsvik et al., 2021). Experimental exposure studies have primarily investigated the effect of MeHg on gene expression in the liver, muscle, and brain tissue of zebrafish. In the liver, the genes affected by Hg exposure were enriched in pathways related to xenobiotic metabolism, oxidative stress, and inflammation (Zhang et al., 2020, Song et al., 2022). In muscle, genes associated with muscle development and function were affected by Hg exposure, while in the brain, Hg exposure was found to impact genes involved in neuronal development, synaptic transmission, neurotransmitter metabolism, and calcium transporters (Gonzalez et al., 2005, Cambier et al., 2009, Cambier et al., 2010, Ung et al., 2010., Zhang et al., 2016, Zhang et al., 2020). One study in zebrafish also identified genes encoding calcium transporters that trigger immune response activation (Cambier et al., 2012) and the effects of various factors on calcium signalling pathways, nuclear receptors, and ion channels (Ung et al., 2010, Rico et al., 2011, Biswas and Bellare, 2021). At the proteomic level, Hg exposure has been found to be associated with gap junction signalling, oxidative phosphorylation, and mitochondrial dysfunction (Rasinger et al., 2017). In brief, exposure to mercury (Hg) affects both neuronal signalling and immune responses, all consistent with our findings.

An important aspect of the study of the consequences of pollution is to understand whether biological changes are adaptive and if such adaptation is the result of natural selection or phenotypic plasticity (Maes et al., 2005; Reid et al., 2016). Using the same populations and individuals as this study, a genome-wide association study by Calboli et al. (2021) identified various candidate genes, including cellular metal binding processes and zinc ion binding, that may play a role in mitigating the effects of Hg accumulation in muscle tissue. The data suggests that there is a genetic basis of adaptation to Hg pollution. None of the genes identified in the present study overlapped with the genes suggested by Calboli et al. (2021). Such overlapping genes were not expected, given our focus on gene expression in the spleen, rather than in muscle tissue. Yet, a lack of overlap may also indicate other adaptive or non-adaptive mechanisms underlying the biological responses to Hg pollution, or that the physiological plasticity in gene expression partially offsets the effect of long-term Hg-induced selection in sticklebacks. Wild populations of fish exhibit gene plasticity in response to ecotoxicological factors such as tolerance and adaptation in selection (Reid et al., 2017). Understanding how organisms respond to environmental stressors can also help us develop strategies to mitigate the negative effects of pollution on ecosystems and protect biodiversity (Hamilton et al., 2016). For instance, genetic diversity may have provided enough standing genetic variation to killifish populations in polluted environments, enabling them to adapt quickly to changing environmental conditions (Reid et al., 2017). In contrast, populations with lower genetic diversity may be less able to adapt and are at greater risk of extinction (Reid et al., 2016).

Governmental agencies monitor Hg concentration in fish to assess health risks, as they often exceed the reference concentrations of the FAO/WHO and European Commission (Capodiferro et al., 2022, Baeyens et al., 2003, Kerambrun et al., 2013). The average Hg concentration in muscle tissue of the stickleback individuals selected for this study ranged from 22.0 to 326 ng g^−1^ dry weight (Calboli et al., 2021). Assuming a typical moisture content of 80 % in fish tissue, these values correspond to 4 to 65 ng/g wet weight. While none of the populations in this study exceeded the FAO/WHO guidelines for human consumption (500ng/g wet weight; Condiniet et al., 2023), the three populations with high mean mercury accumulation did exceed the European Biota Quality Standard 20 ng/g wet weight (Ribeiro et al., 2015, <u>European Commission, 2013</u>). So, while three-spined stickleback is not consumed by humans, the high values at some locations likely biomagnify in its predators like pike *Esox lucius* and brown trout *Salmo trutta*, which are popular among anglers, as well as piscivorous birds such as kingfisher *Alcedo atthis* and spoonbill *Platalea leucorodia*. Our study, therefore, characterises a molecular response between populations to an environmental threat which can cascade into the food chain.

### Conclusion

RNA sequencing of the spleen tissue revealed 136 differentially expressed genes among three-spined stickleback populations with high and low mean mercury content. The majority of the genes were upregulated in the high mercury group. The detection of differentially expressed genes and their involvement in neuronal signalling and immune responses opens the opportunity for a better understanding of the genetic and biological characteristics of organisms living in mercury-polluted environments. Such knowledge contributes to the management of aquatic ecosystems and the protection of biodiversity for current and future generations. Strategies for further research encompass controlled experiments allowing to determine biological responses specific to mercury exposure, as well as field studies linking gene expression to a broader range of multiple environmental stressors.

## Acknowledgements

The Research Foundation-Flanders (FWO) supported our research with a grant to FAMV and GDB entitled “Adaptive responses of an aquatic vertebrate to chemical pollution - KWIKSTICK” (grant G053317N). We thank Leona Milec for her insightful discussions throughout the research project, Martina Kopp for guidance with RNA sequencing, and Kanchana Bandara and Prabhugouda Siriyappagouder for their help with Figure 1A.

## Author contributions

Conceptualization – Brijesh S. Yadav and Joost A.M. Raeymaekers;

Formal analysis – Brijesh Singh Yadav; Aruna M Shankregowda and Joost A.M. Raeymaekers;

Investigation – Brijesh S. Yadav, Aruna M Shankregowda, Vyshal Delahaut, Federico C. F. Calboli, Deepti M. Patel, Marijn Kuizenga, Lieven Bervoets, Filip A.M. Volckaert, Gudrun De Boeck, Fabien C. Lamaze and Joost A.M. Raeymaekers.

Methodology – Brijesh S. Yadav, Aruna M Shankregowda, Fabien C. Lamaze and Joost A.M. Raeymaekers;

Supervision – Joost A.M. Raeymaekers and Fabien C. Lamaze;

Writing – original draft – Brijesh S. Yadav, Joost A.M. Raeymaekers and Fabien C. Lamaze;

Writing – review & editing – Brijesh S. Yadav, Aruna M Shankregowda, Vyshal Delahaut, Federico C. F. Calboli, Deepti M. Patel, Marijn Kuizenga, Lieven Bervoets, Filip A.M. Volckaert, Gudrun De Boeck, Fabien C. Lamaze and Joost A.M. Raeymaekers.

## Data availability Online

### Supplement

#### Funding

This study received funding from the post-Doc budget no. (223000-182) in support of FBA, Nord University, Norway

## Supplementary Figures and Tables

**Supplementary Figure 1.**
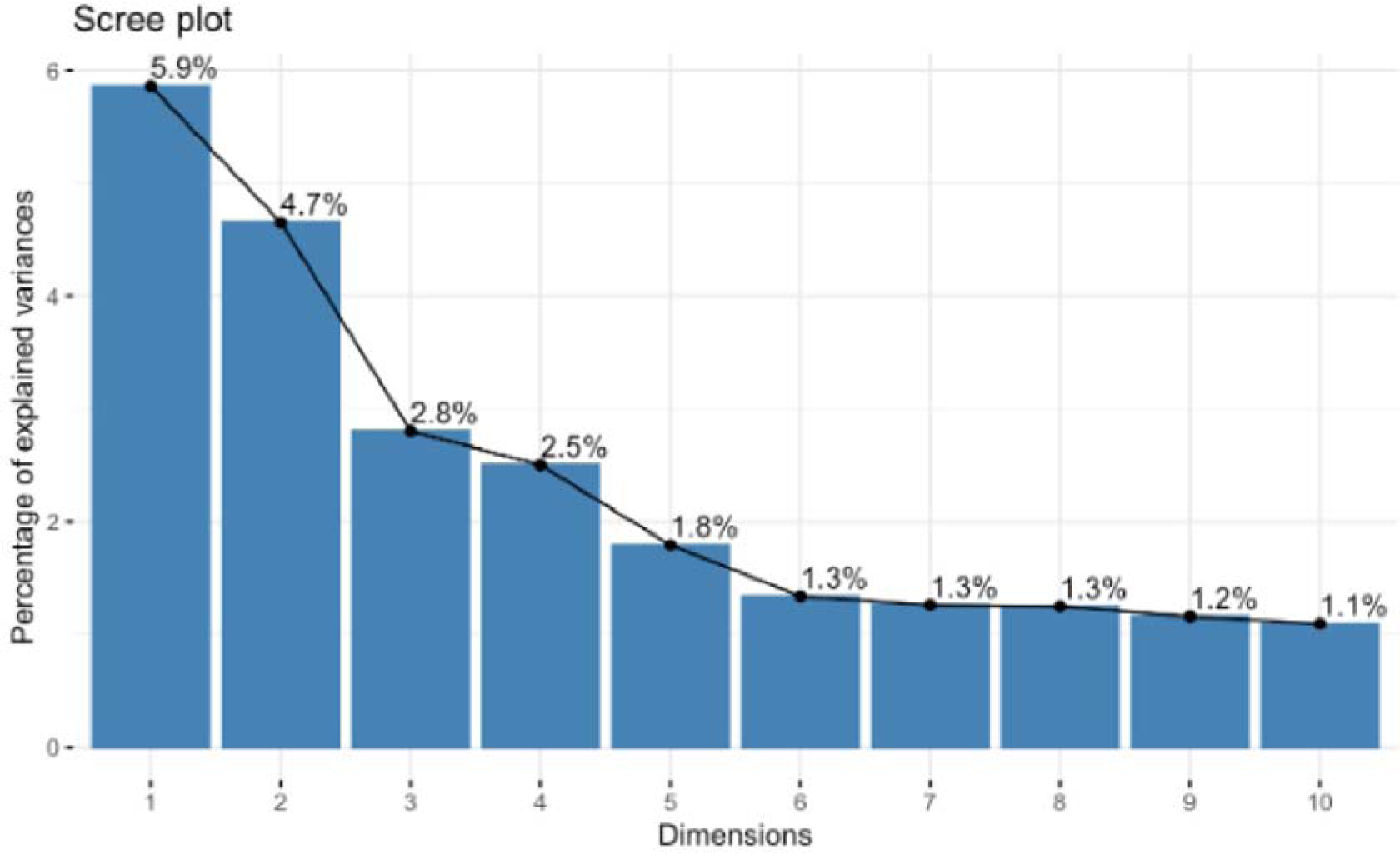
Scree plot of the Principal Component Analysis (PCA) on 28,450 SNP genotypes across 512 individuals.

**Supplementary Figure 2.**
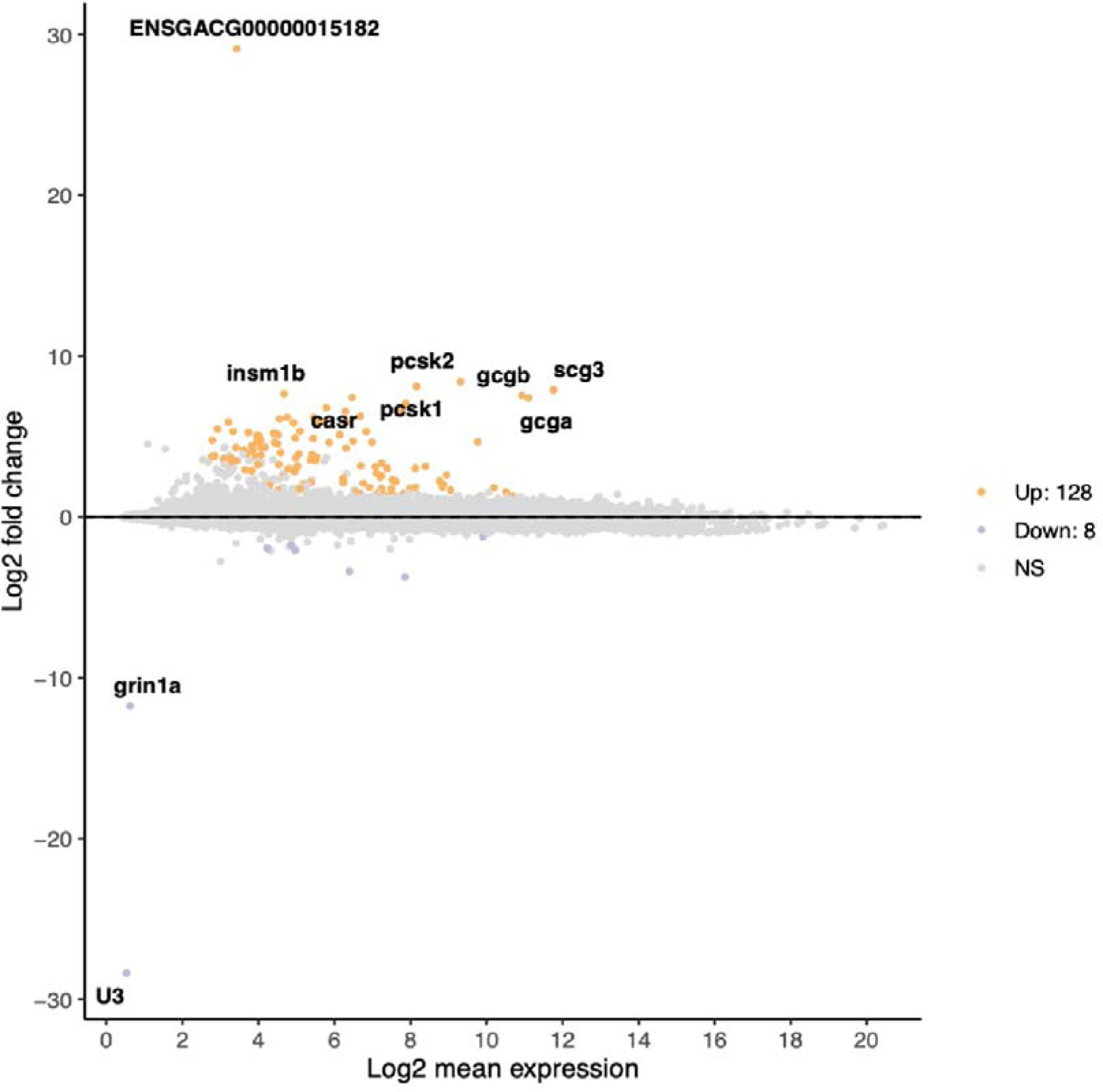
MA plot of the differentially expressed genes among individuals from the high Hg and low Hg group. The plot marks the differentially expressed genes with an adjusted s-value below 0.1 and |Log2 fold change| ≥ 1.

**Supplementary Figure 3.**
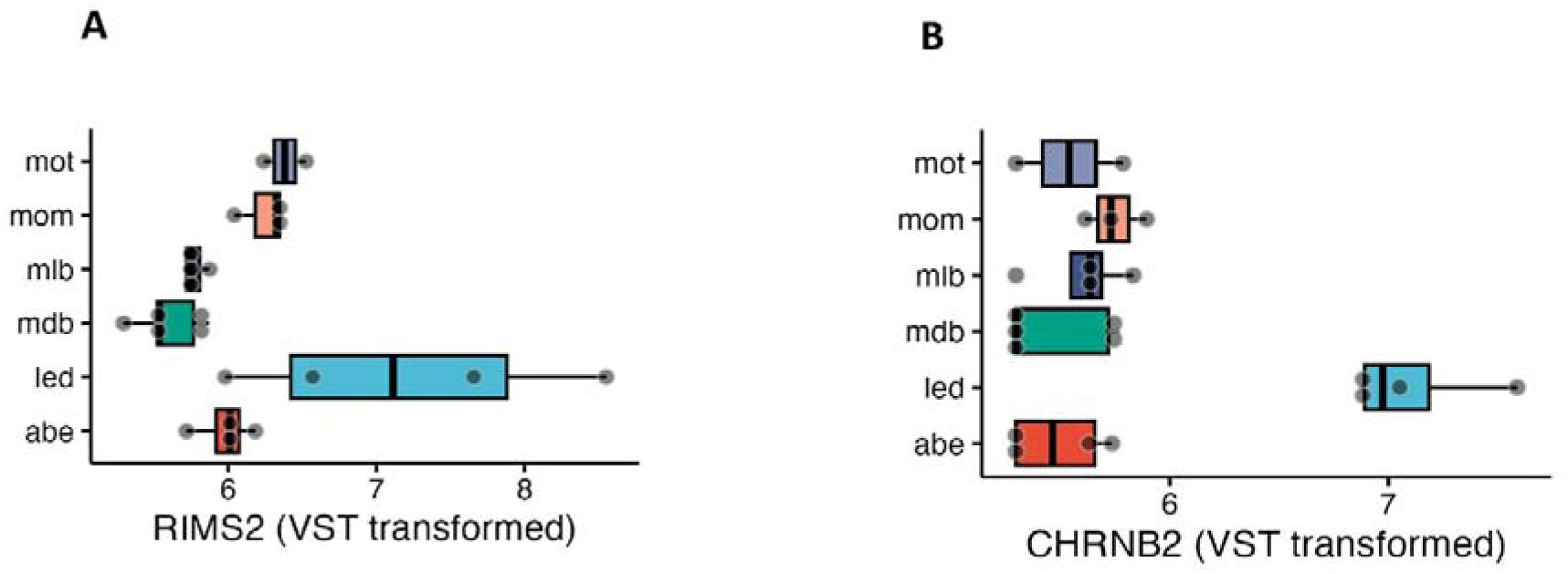
Boxplots of differential expression by location for two genes involved in neuronal functions, (A) RIMS2 and (B) CHRNB2.

**Supplementary Figure 4.**
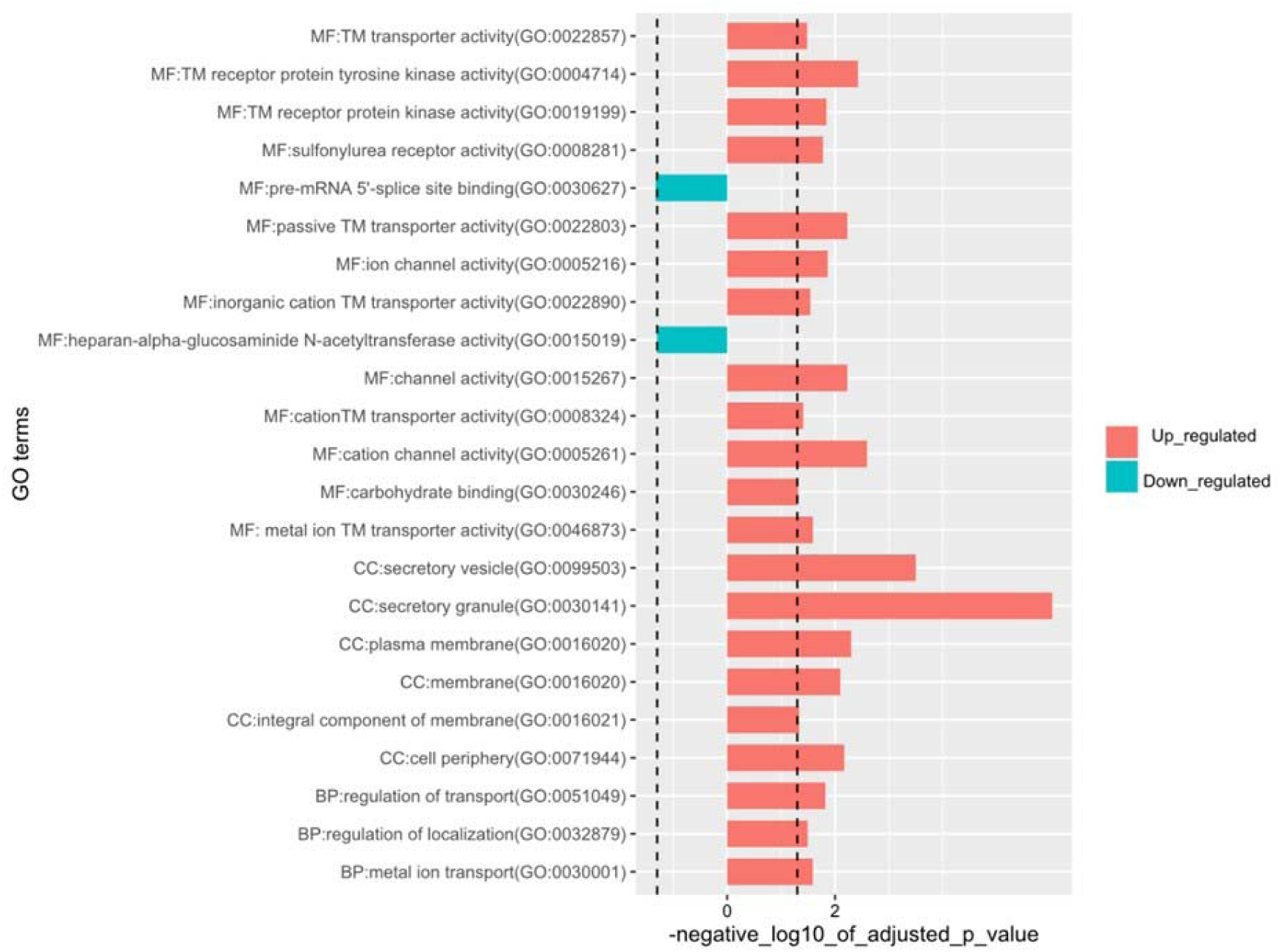
Gene ontology analysis of candidate genes based on their associated biological processes (BP), molecular functions (MF), and cellular components (CC). The significance of each GO term is represented through the -Log_10_ transformation of the g:SCS adjusted p-values, with a higher -Log_10_(P_g:SCS_) indicating a smaller adjusted p-value and thus greater significance.

**Supplementary Figure 5.**
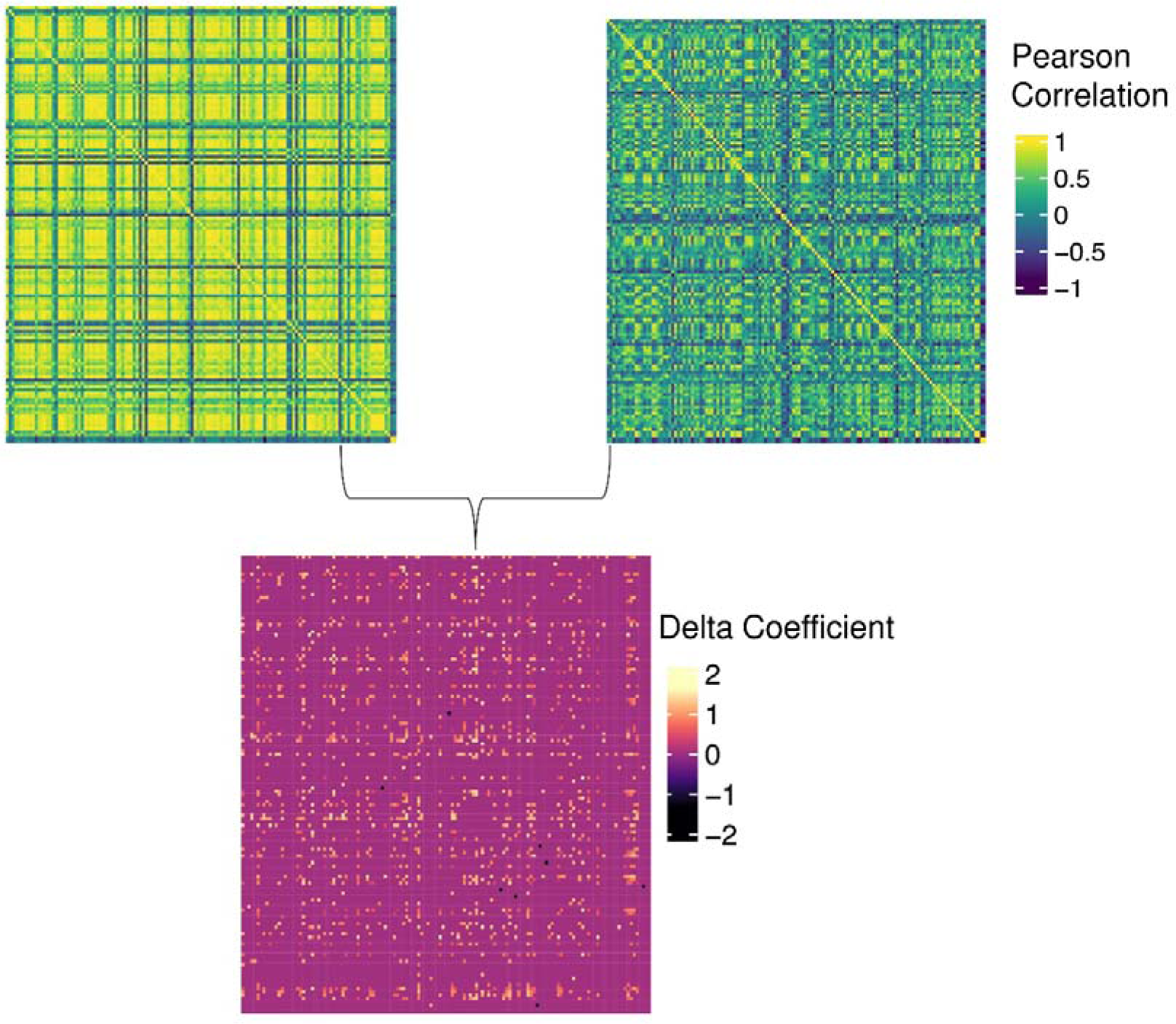
Upper left and upper right panel: correlation matrices of the expression levels of the genes that are differentially expressed in populations from the high Hg (left) and low Hg (right) group. Lower panel: difference (delta) between the high and low correlation matrix. Only significantly dysregulated hub genes at an FDR < 0.01 are shown (non-zero values).

**Supplementary Table 1.**
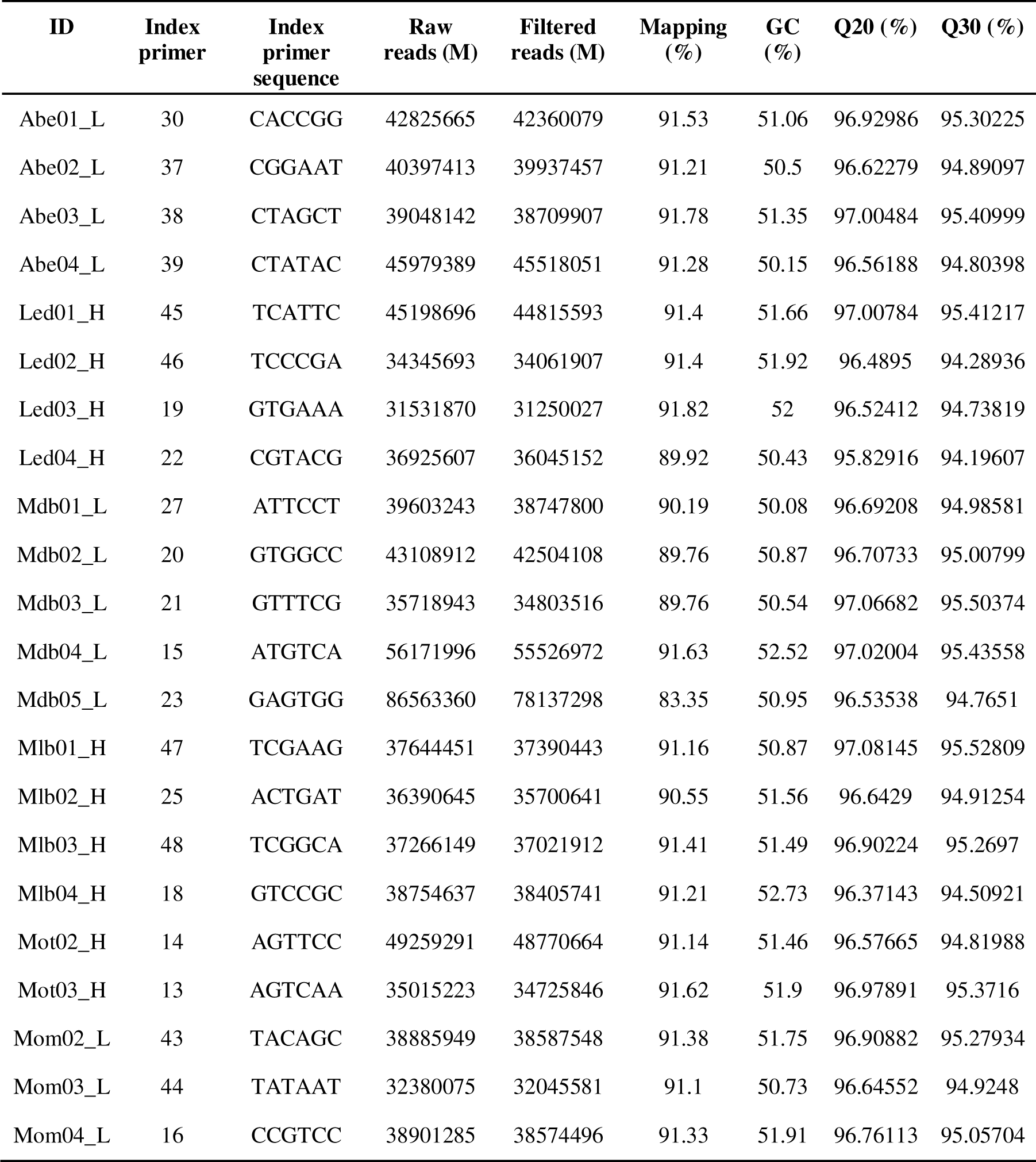
Detailed sequencing output of the spleen transcriptome of the 22 three-spined sticklebacks. ID’s with suffix “H” and “L” mark individuals from the high (N = 10) and low Hg group (N = 12), respectively.

**Supplementary Table 2.**
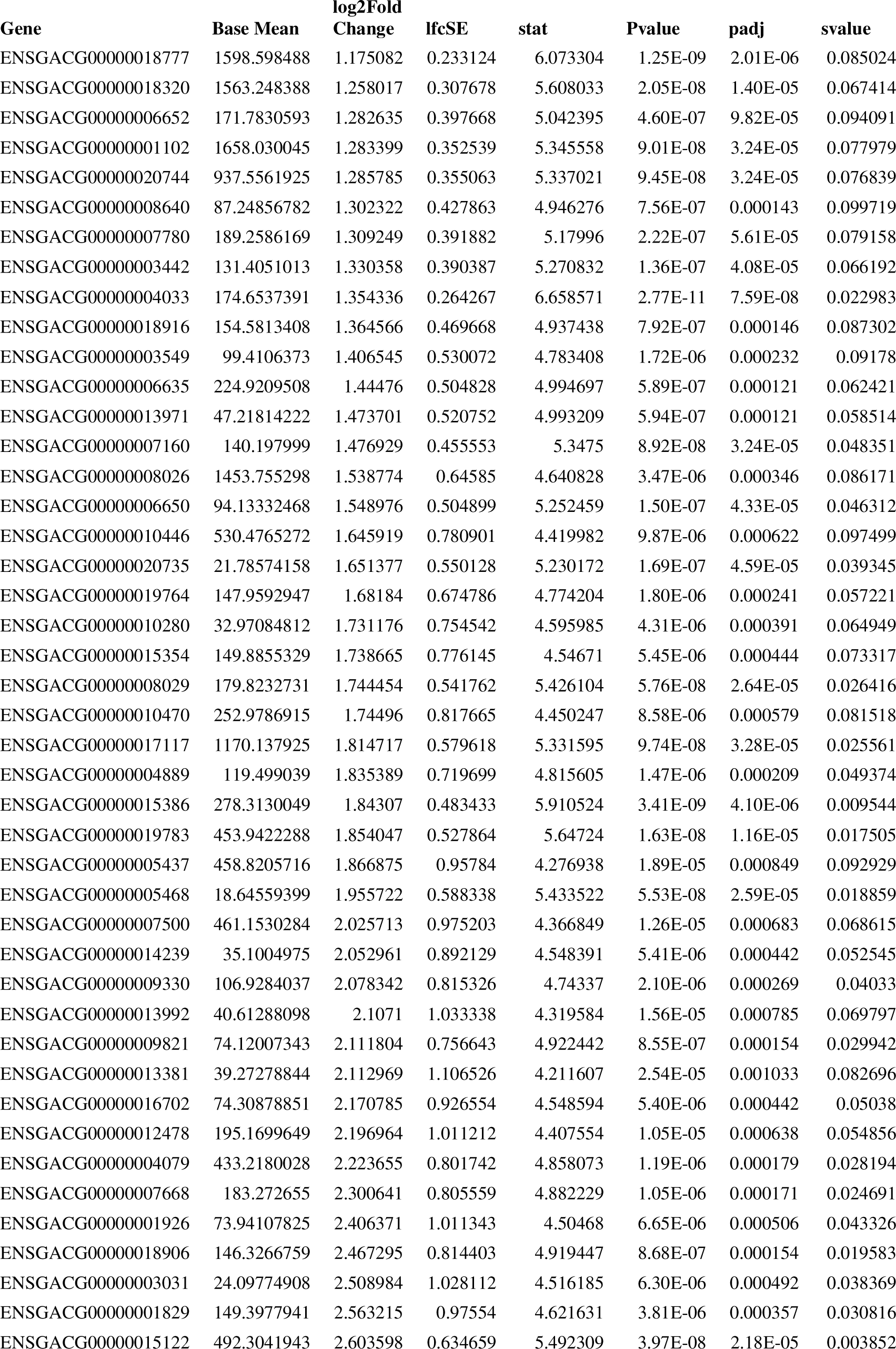
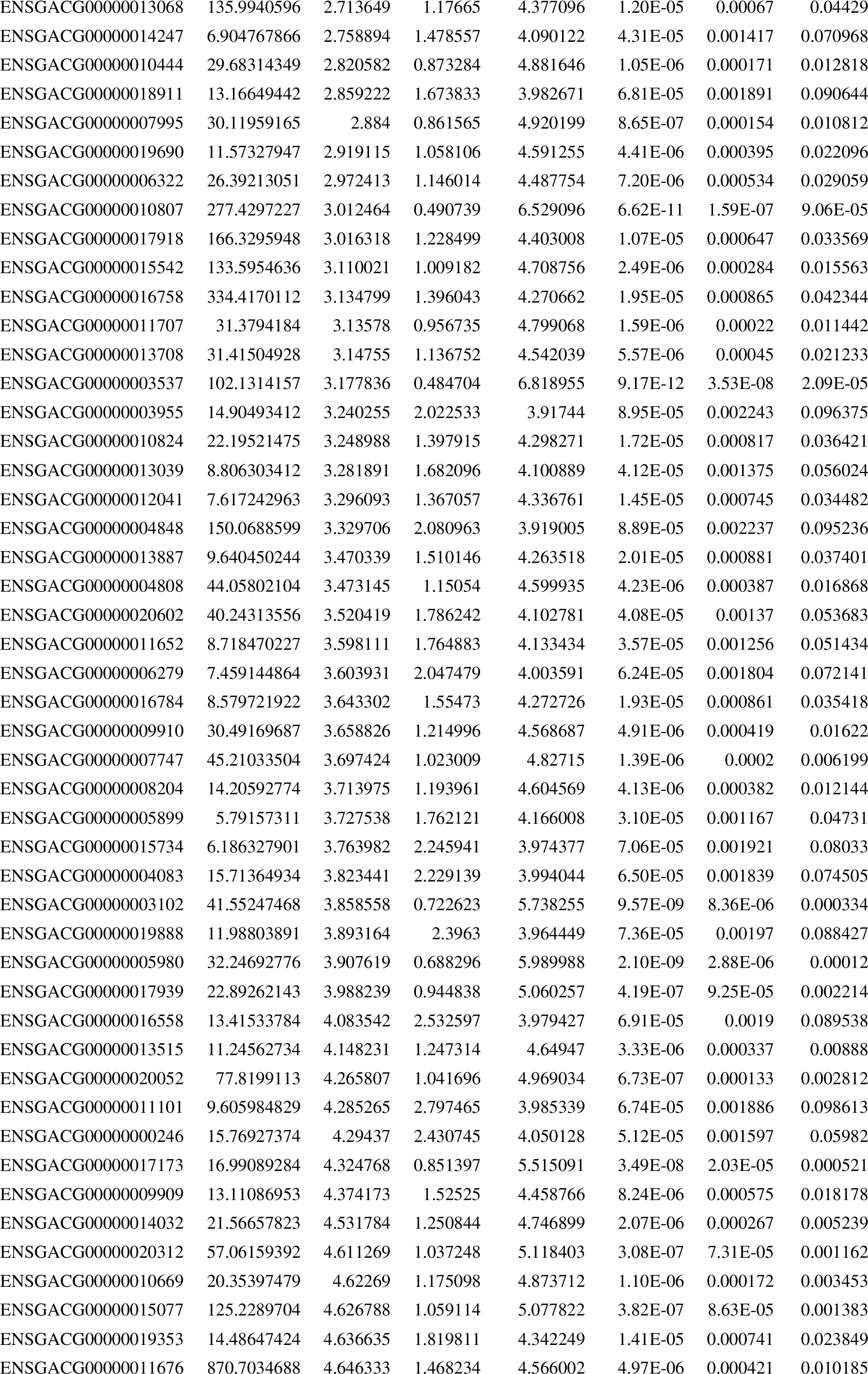
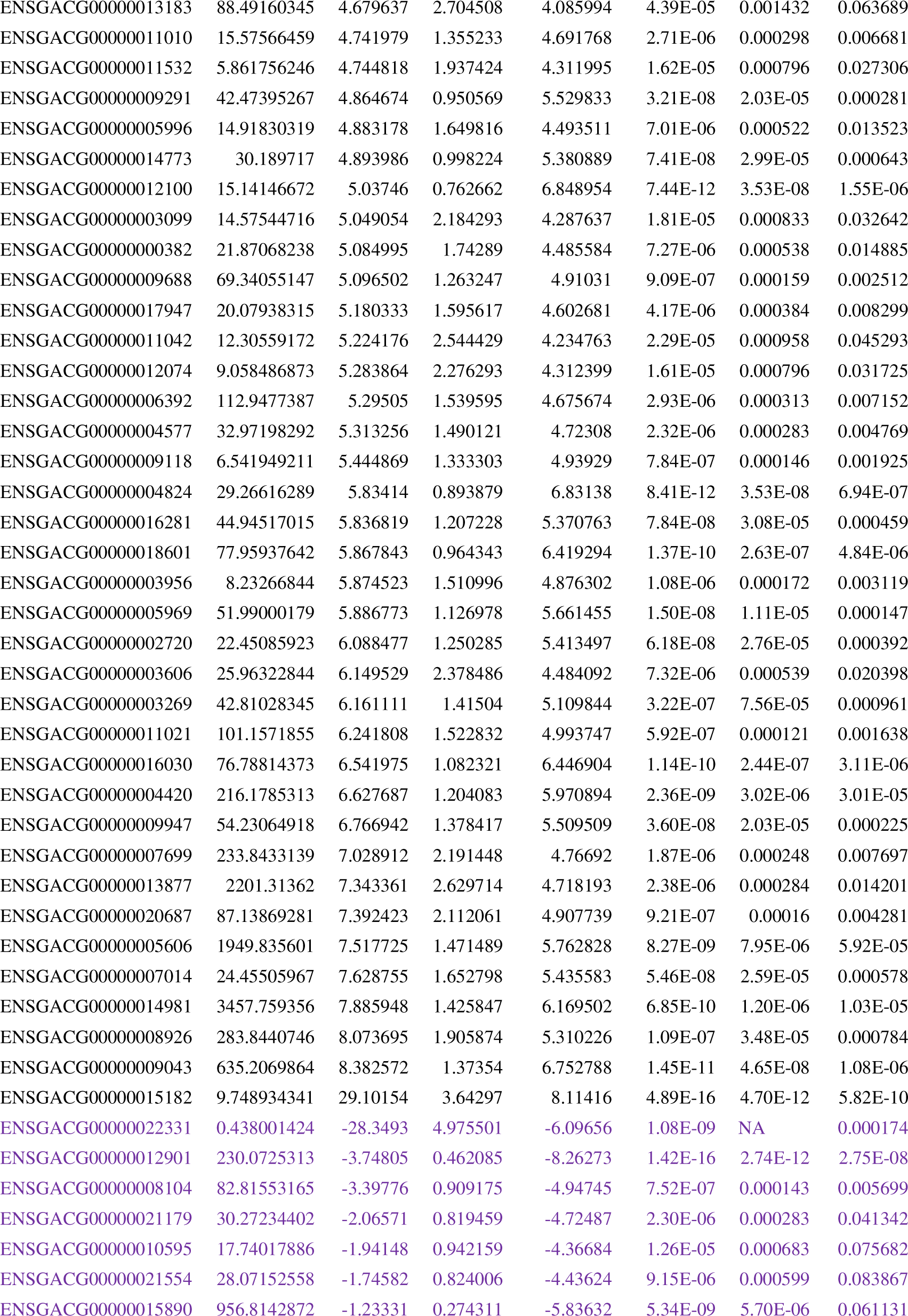
The differentially expressed genes from the high and low Hg environment with an adjusted s-value below 0.1 and |Log2 fold change| ≥1.

